# Chytridiomycosis infection and heat compromises sperm quality in a threatened frog

**DOI:** 10.64898/2026.01.26.701870

**Authors:** Rose Upton, Anne Ibbotson, Kaya Klop-toker, Lachlan Campbell, Nadine Nolan, Phil Jobling, Michael Mahony, John Clulow, Natalie Calatayud, Alex Callen

**Affiliations:** Amphibian Integrated Conservation Unit, Centre for Conservation Science, School of Environmental and Life Sciences, The University of Newcastle, Australia, Callaghan NSW 2308; School of Biomedical Sciences and Pharmacy, The University of Newcastle, Australia, Callaghan NSW 2308; Ian Potter Australian Wildlife Biobank, Melbourne Museum, Carlton, Victoria 3053, Australia

**Keywords:** phenotypic plasticity, terminal investment, reproductive–immune trade-off, thermal stress, spermatogenesis

## Abstract

Environmental change is reshaping wildlife reproduction through increasing temperatures and the spread of emerging infectious diseases, yet the physiological consequences of managing these stressors remain poorly understood. Amphibians are particularly vulnerable due to their ectothermy and high susceptibility to chytridiomycosis, caused by *Batrachochytrium dendrobatidis* (*Bd*). Here, we examine how *Bd* infection and thermal treatment interact to influence sperm quality and reproductive investment in male green and golden bell frogs (*Ranoidea aurea*), a species that has suffered severe population declines. Moderate *Bd* infection was associated with elevated sperm concentration relative to uninfected and heavily infected males, consistent with increased short-term reproductive investment under elevated mortality risk. However, severe infection led to pronounced reductions in sperm concentration and motility. Thermal treatment successfully eliminated *Bd* infection but imposed substantial reproductive costs: sperm concentration declined following treatment and remained significantly reduced six months later, despite partial recovery of sperm motility and membrane integrity. These results indicate persistent impairment of spermatogenic capacity rather than transient suppression. Our findings reveal that disease and thermal stress jointly shape amphibian reproductive outcomes through context-dependent trade-offs between immune defence and gamete production. While mild infection may trigger short-lived increases in reproductive output, both severe infection and pathogen clearance via thermal exposure impose lasting constraints on fertility. These results highlight an underappreciated cost of disease mitigation and suggest that increasing thermal extremes associated with climate change may further limit amphibian reproductive resilience, with important implications for conservation management and population persistence.

## 1. Introduction

Environmental change is reshaping wildlife health and reproduction through shifts in temperature, altered precipitation, and the spread of emerging diseases. These stressors disrupt endocrine and immune systems, leading to reduced fertility, abnormal development, and increased disease susceptibility across vertebrates (1). Global assessments show that amphibians are among the most affected, with disease and climate change together driving the majority of recent declines (2). As ectotherms, amphibians depend heavily on environmental temperature and moisture to regulate physiological function, making them particularly vulnerable to rapid climate variability. Recent analyses indicate that global warming is already pushing several species beyond their physiological heat limits, even within shaded or aquatic habitats that were once thermally stable (3). These findings highlight how increasing temperature extremes and disease outbreaks act together to constrain amphibian survival in a rapidly changing world (4).

The ability of vertebrates to cope with such instability depends on phenotypic plasticity, the capacity to adjust physiological and behavioural traits to maintain homeostasis and reproductive function under changing conditions (5). In amphibians, this plasticity is central to balancing immune defence, breeding effort, and energetic costs across fluctuating environments. Yet, the limits of plasticity become apparent when environmental change outpaces physiological compensation, as occurs under combined heat and disease stress. In such cases, reproductive and immune processes can become misaligned, resulting in reduced gamete production, impaired fertility, and compromised resilience (6, 7). As outlined in the *Amphibian Conservation Action Plan*, these physiological thresholds and the limited capacity of many amphibians to adapt or shift their ranges underscore the urgency of understanding reproductive resilience under compounding stressors (4).

Among the most devastating examples of how disease and climate interact to threaten biodiversity is the emergence and persistence of chytridiomycosis, caused by the fungal pathogen *Batrachochytrium dendrobatidis* (*Bd*). This pathogen has driven the extinction of more than 90 amphibian species globally, including at least seven in Australia, and continues to contribute to the decline of more than 500 others, making it one of the most destructive infectious agents known to science (8-10). The relationship between *Bd* and temperature is complex: infection thrives in cool, moist environments, whereas higher temperatures can suppress the pathogen but impose physiological strain on the host (11, 12). With climate warming predicted to increase the frequency of extreme thermal events, amphibians now face overlapping pressures of thermal stress and persistent infection, each influencing immune competence and reproductive performance (7).

The green and golden bell frog, *Ranoidea aurea* (previously *Litoria aurea*), exemplifies this intersection between disease, physiology, and climate vulnerability. Once widespread along the southeastern coast of Australia, the species has suffered extirpations across more than 90 % of its range over recent decades (13). Chytridiomycosis is considered the primary driver of its decline (14, 15), and few management options exist for wild populations (but see (16)). Because *R. aurea* can be maintained successfully in captivity and responds well to assisted reproductive technologies (17-19), it provides an ideal system for investigating the physiological effects of disease and potential management strategies recently trialed, including vaccination or heat-based infection clearance (20). These management approaches reflect the integrated conservation frameworks promoted by *Amphibian Conservation Action Plan* (4), which emphasize coordination between ex situ population management, disease control, and reproductive technologies.

Conservation strategies have predominantly focused on preventing extinction through the establishment of quarantined, disease-free colonies and through controlled thermal treatments aimed at clearing infection, but there is little information examining the interactions between reproductive plasticity, disease, and thermal tolerance and recovery. Linking these management strategies to their physiological consequences provides essential insight into how amphibians balance infection control with the maintenance of reproductive capacity. Elevated temperatures are known to disrupt spermatogenesis through oxidative stress, mitochondrial dysfunction, and germ-cell apoptosis in many vertebrates (21, 22), and comparable effects are expected in ectotherms. Such cellular stress pathways may underlie the trade-offs observed in amphibians undergoing thermal treatment, where infection clearance can coincide with declines in reproductive capacity (23).

The energetic conflict between immune defence and reproduction provides a theoretical foundation for understanding these trade-offs. The terminal investment hypothesis predicts that individuals facing reduced survival prospects will allocate resources toward immediate reproductive effort rather than future survival (24-26). While this response has been documented across diverse taxa (27, 28), amphibians offer a unique system in which to test how infection and environmental stress interact to shape reproductive investment. Evidence from *Bd*-infected species shows variable outcomes: some populations exhibit increased reproductive effort (12, 29), while others show reduced gonadal investment and sperm quality (30, 31). This variation reflects the context-dependent and plastic nature of reproductive responses to infection severity and thermal environment.

This study examines how *Bd* infection and elevated temperature interact to influence sperm quality and reproductive investment in male *Ranoidea aurea*, a species that has undergone severe declines due to chytridiomycosis. We specifically aimed to: (i) determine how variation in *Bd* infection severity affects sperm quality; (ii) assess the effects of initial infection status and heat-based pathogen clearance on sperm quality; and (iii) quantify the short- and long-term consequences of thermal treatment for sperm quality, including effects persisting six months post-treatment. By explicitly linking infection burden, disease mitigation, and recovery time, this study provides mechanistic insight into how immune activation and thermal stress regulate reproductive trade-offs and constrain amphibian resilience under ongoing climate change.

## 2. Methods

### 2.1 Study species and animal husbandry

All animals in this study were sourced from the University of Newcastle’s *R. aurea* captive breeding colony. Frogs were housed in seminatural outdoor mesocosms (75 cm × 75 cm × 110 cm) containing approximately 25% area terrestrial gravel substrate with vegetation, 75% water and stacked brick microrefuges throughout. Frogs were fed with crickets one to two times per week, supplemented with calcium and vitamin powder (Multical Dust; Vetafarm, Wagga Wagga, NSW, Australia), and exposed to natural day–night light regimes, being able to bask at will. Frogs were transferred to the laboratory for heat treatment from the outdoor holding mesocosms in July 2024 after *Bd* was detected during disease surveillance. Once indoors, frogs were housed in plastic terraria (30 ×18 x 20 cm), with 25% terrestrial environment (autoclaved pebbles) and 75% aquatic environment (aged tap water) and plastic piping as hides.

All frogs underwent a precautionary protocol of heat treatment regardless of quantitative polymerase chain reaction (qPCR) results to ensure no *Bd* infection remained in the colony and opportunistic sampling was performed, as outlined below. Frogs were placed in UV- and temperature-controlled cabinets (TRISL – 1175, Thermoline Scientific Equipment, Wetherill Park, NSW) on a 12:12 UV light cycle started at 25°C. Temperature was increased at 2°C per day until 37°C was reached, where the temperature was held for 6 h. This temperature has been shown to kill *Bd* within 4 h (32). Subsequently, temperature was reduced at 2°C per day until the original 25°C was reached and frogs were reswabbed to confirm disease had been cleared. Frogs were kept at 24-25°C until the conclusion of the experiment in September, 2024.

All captive husbandry and experiments were conducted in accordance with institutional and national Australian standards for the care and welfare of animals, and in accordance with the University of Newcastle’s animal ethics approval (A-2022-207 and A-2024-413). Animals were originally collected under NSW Scientific Licence SL101269.

### 2.2 Opportunistic sampling

While infection of the frogs in this study was not part of a planned experiment, the outbreak event allowed opportunistic sampling to investigate the effects of disease and treatment on sperm quality. On the same day, but prior to commencing heat treatment, all frogs were weighed (to nearest 0.1 g) and snout-vent-length (SVL) and right tibia measured (to the nearest 0.1 mm). All frogs were swabbed in order to determine *Bd* infection rates and loads via quantitative polymerase chain reaction. Swabs were stored at -30°C until DNA extraction. Forty-two frogs in total were in the cohort affected by this outbreak, with six deaths due to *Bd*. On day 2 of heat treatment, 13 mature males were selected for sperm induction, blood collection and biopsy of the toe webbing. All frogs that were selected were without advanced symptoms and had consistent and prompt righting reflexes of under 1 second. In addition, two males that were found deceased at the discovery of the outbreak were dissected, and sperm quality from testicular macerates was assessed, as per section 2.2.1.

#### 2.2.1 Induction of spermiation and sperm collection

Frogs were removed from heat treatment in the morning, prior to increasing temperature for the day, for two hours and kept at 25°C. After urine sample containing sperm (spermic urine) were collected, animals were returned to heat treatment and temperature raised to 27°C. As frogs were originally sourced from wild populations, age was unknown (however males reach sexual maturity at around 1 year of age and all displayed secondary sexual characteristics indicative of sexual maturity (darkened nuptial pads and coloured throat)). The weight range of males used in the study was 21.5 ± 3.8 g (mean and standard error, n = 13). None of the 13 males selected for spermiation experiments died due to infection.

A sample of urine was collected prior to hormone injection and found to be aspermic. Males were injected with 5 IU/g body weight human chorionic gonadotropin. Previous studies have shown no difference in sperm concentrations yielded between 10 and 20 IU/g of body weight and recent data has refined the protocol to use less hormone per injection (Upton, unpublished data). Hormones were diluted in simplified amphibian ringer (SAR; 113 mM NaCl, 1 mM CaCl2, 2 mM KCl, 3.6 mM NaHCO3; ∼220 mOsm/kg; recipe (33)) to a final volume of 200 µl. Hormones were administered subcutaneously via the dorsal lymph sac using a 31-gauge insulin syringe (*BD*, New Jersey, United States). Frogs were placed in individual inflated plastic bags with aged tap water sprayed into the bag for hydration. Half an hour after injection, a thin plastic tubing (Tomcat feline catheter; Argyle), was used as a catheter to collect spermic urine samples. The tip was gently placed into the cloaca and gently moved in and out to facilitate urine collection by capillary action. Spermic urine was collected at intervals of 10-15 minutes over an hour period and pooled in 1.5 mL Eppendorf tube for analysis. Samples were kept on ice until volume, motility and sperm concentration could be determined (within the same day). Motility analyses were performed immediately after collection to reduce possible effects of time post-collection, followed by assessment of membrane integrity and sperm concentration.

In two frogs found recently deceased (not included in sperm induction experiment), both testes were removed and placed in 1 mL SAR. A sperm suspension was prepared by gently teasing apart testicular tissue with fine forceps to release sperm. Sperm quality assessments, described below, were conducted immediately after maceration.

#### 2.2.3 Post-infection sampling

Sperm sampling was repeated five weeks following the conclusion of heat treatment (seven weeks following initial outbreak detection), in September 2024. While the duration of spermatogenesis is unknown in this species, previous work has shown the duration of spermatogenesis to be approximately 40 days in *Lithobates catebeianus* (34). Thus, to determine whether any short-term effects of *Bd* infection were still present, this period was chosen to allow a full cycle of spermatogenesis to occur post-infection. As results indicated a possible effect of heat treatment on sperm quality, a subset of four of the 13 frogs were sampled again six months following completion of heat treatment. The full cohort was unable to be sampled at six months post-heat treatment due to frogs entering breeding protocols. All four frogs sampled at 6 months were positive for *Bd* at their initial timepoint but were cleared of *Bd* by the initial heat treatment.

### 2.3 DNA extraction and quantitative Polymerase Chain Reaction

A standardised swabbing technique was employed over the epidermal surfaces prone to high infection loads, which involved wiping both sides of the ventral skin 8 times, the inner and outer thighs 4 times and the hind and fore feet 2 times each (35). DNA was extracted from swab tips and quantified following standard qPCR Taqman assay methods (36), using a Biorad CFX96 system. Each swab was assessed for the presence of *B. dendrobatidis* in triplicate using a 1/10 diluted sample from the DNA extracts. Standard curves were generated using commercially prepared standards (Pisces Molecular. Boulder, CO, USA) and were serially diluted to prepare five known *B. dendrobatidis* concentrations (18700, 1870, 187, 18.7 and 1.87 genomic equivalents (GE)/⍰l). Final data were expressed as mean genomic equivalents per microlitre of a standardised extract volume (5⍰µl) from three replicates of the same swab sample. A frog was considered uninfected if the mean swab result was <1.00 zoospore equivalents (37), if at least two of the triplicates did not amplify (23, 35) or where amplification occurred >40 cycles, which was interpreted as non-specific binding. Where at least two triplicates amplified, the average of all three wells was used if no presence of reaction inhibition was present. Inhibition was detected with inclusion of an internal positive control (Taqman Exogenous Internal Positive Control Reagents; Applied Biosystems, Massachusetts, United States) in a single replicate of each sample. Where the internal control did not amplify, samples were considered inhibited and rerun at a 1:100 dilution. Data were multiplied by either 10, or 100, to account for the dilution step during the extraction process.

### 2.4 Sperm Quality assessments

Concentration (cells/mL) of sperm in urine samples were assessed with an Improved Neubauer haemocytometer counting chamber at each collection interval. Sperm were diluted, as necessary, with 1.7% NaCl and the dilution factor was accounted for in the final calculation. Ten microlitres of sperm solution was pipetted into each of two chambers, and the number of sperm in at least five secondary quadrats (per chamber) was counted, such that at least 80-100 sperm cell were counted per sample, and used to calculate total sperm per millilitre. The number of secondary quadrats was accounted for in the final calculation.

Due to high concentrations of spermic urine present in samples, motility was analysed in all samples after a 1:5 dilution in reverse osmosis (RO) water. While analysis of undiluted samples was preferable, high densities of sperm made visualization of individual sperm cells impossible. Sperm were categorised as either non-motile, motile without forward-progression or motile with forward progression, and at least 100 sperm were counted at 400x magnification under phase contrast optics. Proportions were calculated for forward-progressive and total motility (sperm motile with or without forward-progression). Membrane integrity was determined using a dye-exclusion assay. Eosin-Y (Sigma Aldrich; E4009; 0.4% w/v diluted in 0.9% w/v saline) was mixed 1:1 with sperm samples, incubated for at least 30 s at room temperature and then scored under a coverslip using brightfield microscope at a magnification of 400x. Sperm that stained pink were scored as compromised, whereas sperm with a clear cytoplasm were scored membrane intact. For each sample, 100–200 sperm were scored per count, and two counts conducted per sample. If two counts from the same sample differed substantially, a third count was made and included in the analysis.

### 2.5 Statistics and experimental design

Generalised linear mixed models (GLMM) were used to fit Poisson and binary logistic regressions to four sperm parameters: sperm concentration (cells/mL), proportion forward-progressive motility, proportion total motile, and proportion membrane intact. For sperm concentration, raw counts were used with an offset of the log reciprocal of an adjustment factor (based on the formula for calculating sperm concentration from a hemocytometer, accounting for: dilution factor, volume and the number of quadrats counted) to convert the raw counts to cells/mL. For the motility and membrane integrity measures, the models were interpreted as the proportion of successful cases; i.e. live spermatozoa or motile sperm with the total number of spermatozoa used as weights in the model.

*For Experiment 1: Effect of Bd severity on sperm quality, Bd* infection severity was used as a main effect, where the categories were negative, 0 GE/µL (n=4); moderate, 900-6499 GE/µl (n=5); and high, >6500 GE/µl (n=4). For *Experiment 2: Effect of initial Bd status on sperm quality pre- and post-treatment*, a two-factor design was implemented with *Bd* status prior to heat treatment (negative, n=4; positive, n=9) and time in relation to heat treatment (pre- and post-heat treatment), used as main effects. The interaction between initial *Bd* status and time in relation to heat treatment was not significant and was dropped from the model. For *Experiment 3: Effect of heat treatment on sperm quality*, the time in relation to heat treatment was used as a main effect (pre-, post- and six months post-heat treatment) using a subset of four frogs from each time point. For all models, overdispersion was addressed using animal body condition (weight/SVL) as a random effect. Significance was determined by pairwise comparisons to determine ratios of expected counts between conditions and 95% confidence intervals. Modelled estimated marginal means (EMM) and 95% confidence intervals (CIS) were calculated.

All analyses were completed using R (Version 3.6.2) with R packages glmmTMB used for all GLMM modelling (38). The package emmeans was used to model estimated marginal means (EMMs) and 95% confidence limits were calculated and back transformed to sperm concentrations and motility and membrane intact proportions. Ratios (for sperm concentration data) and odds ratios (for proportion data) comparing treatments were also generated using the package emmeans (39). All graphing was completed using ggplot2 and gridExtra (40, 41).

## 3. Results

### 3.1 Bd load analysis

Four out of thirteen males in this study tested negative for *Bd*. The nine remaining frogs had moderate to high loads of *Bd*, ranging 939 – 20,000 GE/µl and averaging 7815 GE/µl. Two deceased frogs from this study had 403,370 and 34,000 zoospore equivalents respectively. Following heat-treatment, all frogs tested negative for *Bd*.

### 3.2 Experiment 1: Effect of Bd severity on sperm quality

There was a significant effect of *Bd* severity on sperm concentration (likelihood ratio test (LRT) χ^2^(2) = 9.4, *P* = 0.009), with frogs in the moderate infection category having higher sperm concentrations than high load infections and frogs negative for infection (Fig 1a). Ratios and 95% confidence intervals were generated to further compare the differences in sperm concentration in response to *Bd* infection, with larger ratios indicating bigger differences in sperm concentration. Estimated marginal means (EMM) ranged from 4.4×10^8^ to 8.2×10^8^ cells/mL across all categories of severity. Frogs with moderate *Bd* infection had the highest sperm concentration (EMM: 8.2×10^8^, 95% confidence intervals [CIs]: 6.6×10^8^, 1.0×10^9^). Frogs with moderate infection had significantly higher concentrations than frogs negative for *Bd* (Ratio: 1.9, 95% CIs: 1.3, 2.6), with negative frogs having the lowest sperm concentration (EMM: 4.4×10^8^, 95% CIs: 3.4×10^8^, 5.6×10^8^). Frogs with moderate infection also had a significantly higher sperm concentration than frogs with high infection status with an EMM of 6.1×10^8^ cells/mL (95% CIs: 4.7×10^8^, 7.8×10^8^) and ratio of 1.4 (95% CIs: 1.0, 1.9). Frogs with high infection had significantly higher sperm concentration than frogs with no infection (Ratio: 1.4, 95% CIs: 1.0, 2.0).

**Figure 1.**
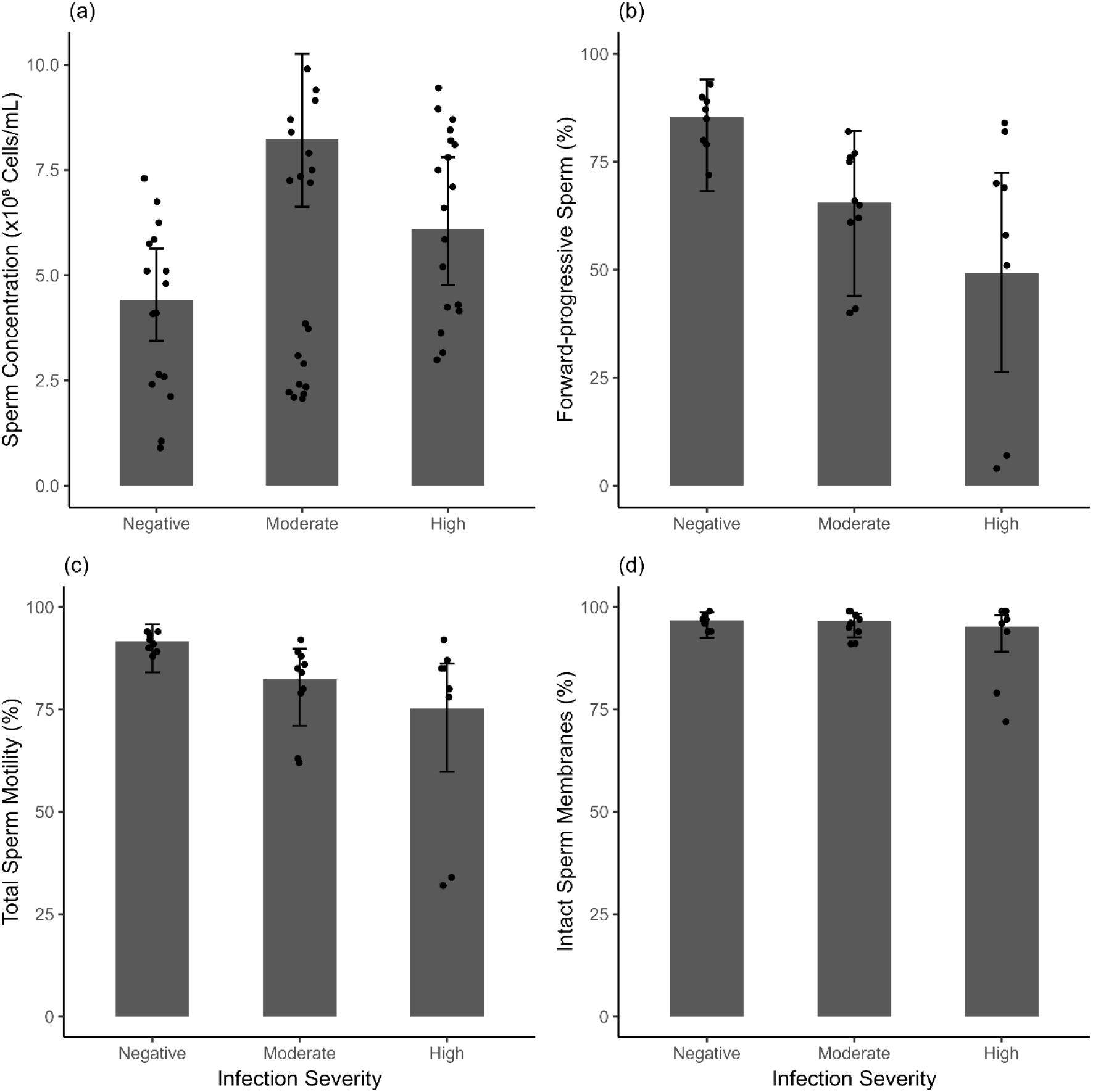
Effect of *Bd* infection severity on induction of spermiation in *Ranoidea aurea* (*n*⍰=⍰13) for (a) sperm concentration (cells/mL); (b) forward progressive sperm (%); (c) total sperm motile (%), and; (d) intact membranes (%). Columns represent estimated marginal means, black dots represent raw data and error bars equal 95% confidence intervals.

There was an effect of *Bd* severity on sperm forward progressive motility (LRT χ^2^(2) = 5.2, *P* = 0.07) and total motility (LRT χ^2^(2) = 5.1, *P* = 0.08) with decreasing motility as infection severity increases (Fig 1b and c). Odds ratios and 95% confidence intervals were generated to further compare the differences in sperm quality. Frogs negative for *Bd* infection had highest sperm motility with 85.4% (95% CIs: 68.2, 94.1) forward progressive motility and 91.7% (95% CIs: 84.0, 95.8) total motility. Frogs with moderate infection had 65.5% (95% CIs: 43.9, 82.2) sperm with forward progressive motility and 82.3% (95% CIs: 71.0, 89.8) sperm with any motility. Frogs with high infection had the lowest motility with 49.2% (95% CIs: 26.3, 72.5) sperm with forward progressive motility and 75.3% (95% CIs: 59.8, 86.2) sperm with any motility. Sperm from frogs with negative infections had significantly higher motility than frogs with high infection loads, with sperm from negative frogs having sixfold higher forward progressive motility (Odds Ratio (OR): 6.0, 95% CIs: 1.5, 24.7) and almost fourfold total motility (OR: 3.6, 95% CIs: 1.3, 10.1). There was approximately a threefold (OR: 3.1, 95% CIs: 0.8, 11.7) and twofold difference (OR: 2.4, 95% CIs: 0.9, 6.3) in forward progressive motility and total motility in negative frogs compared to frogs with moderate infection loads, however this difference was not statistically significant. Frogs with moderate infection loads had higher likelihood of forward progressive motility (OR: 2.0, 95% CIs: 0.5, 7.5) and total motility (OR: 1.5, 95% CIs: 0.6, 4.0) than in frogs with high infection loads, however this difference was also not statistically different

There was no significant effect of *Bd* severity on sperm membrane integrity (LRT χ^2^(2) = 0.4, *P* = 0.8; Fig 1d). Frogs negative for *Bd* infection had highest membrane integrity EMM of 96.8% (95% CIs: 92.5, 98.7), followed by frogs with moderate infection load (EMM: 96.6%, 95% CIs: 92.6, 98.4) and frogs with high infection load (EMM: 95.3%, 95% CIs: 89.1, 98.0).

### 3.3 Experiment 2: Effect of initial Bd status on sperm quality pre- and post-treatment

There was a significant effect of initial infection status and heat treatment on sperm concentration (LRT χ^2^(1) = 3.6, *P* = 0.056 and LRT χ^2^(1) = 7.3, *P* = 0.007 respectively) after five weeks (Fig 2a), with the highest sperm concentration in frogs initially testing positive for *Bd*, pre-heat treatment. Both pre- and post-heat treatment, frogs which initially tested positive for *Bd* had significantly higher sperm concentration than frogs which tested negative, with the ratio of expected sperm concentrations being 1.5 (95% CIs: 1.0, 2.22). Irrespective of initial *Bd* status, sperm concentration was significantly higher prior to heat treatment than post-heat treatment, with the ratio of expected sperm concentrations being 1.7 (95% CIs: 1.2, 2.5). Sperm concentration for positive frogs was 7.0×10^8^ cells/mL (95% CIs: 5.3×10^8^, 9.35×10^8^) and 4.1×10^8^ cells/mL (95% CIs: 3.1×10^8^, 5.4×10^8^) prior to and post-heat treatment respectively.

**Figure 2.**
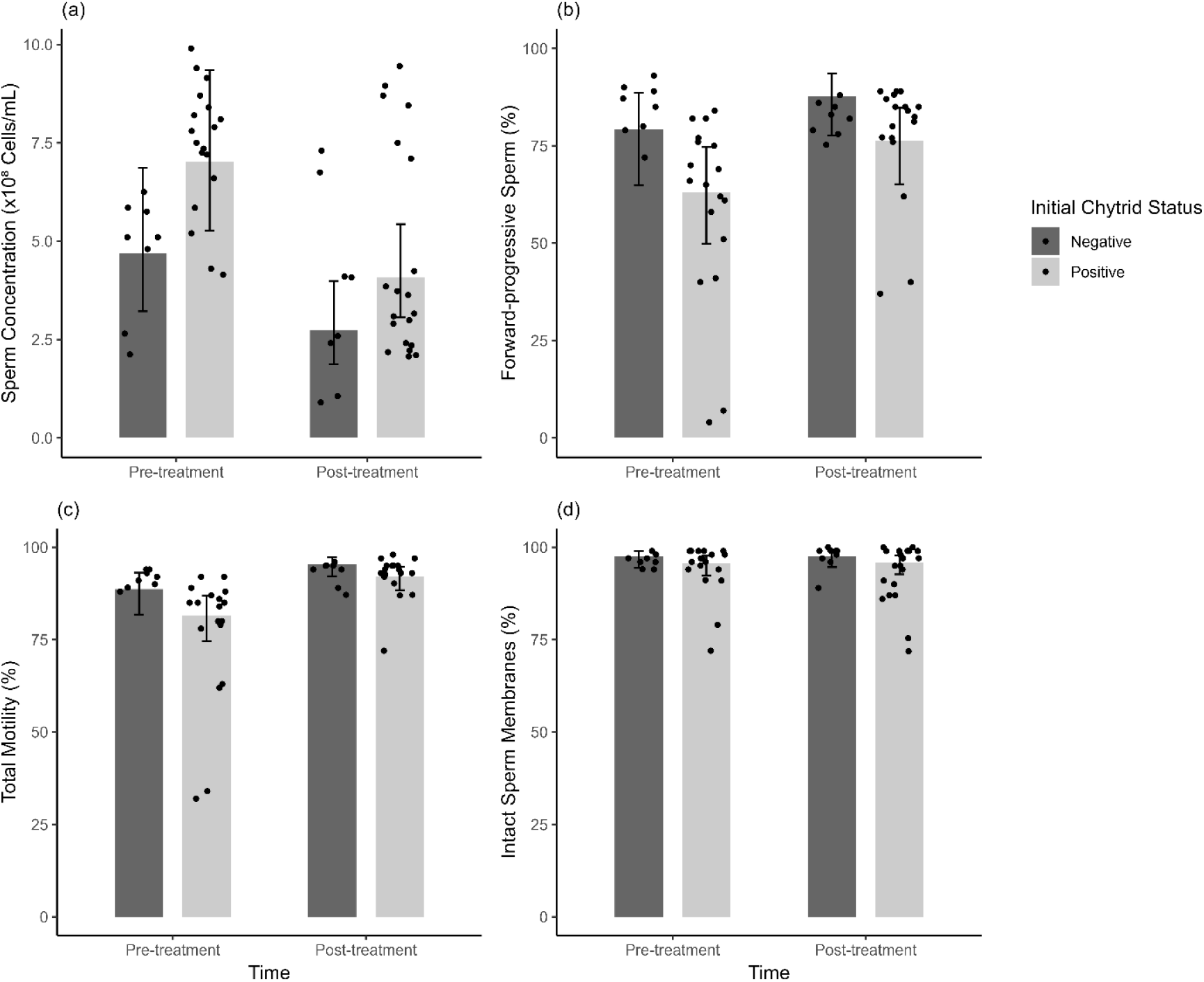
Effect of initial *Bd* infection status and heat-treatment (pre- and post-) on induction of spermiation in *Ranoidea aurea* (*n*⍰=⍰13) for (a) sperm concentration (cells/mL); (b) forward progressive sperm (%); (c) total sperm motile (%), and; (d) intact membranes (%). Columns represent estimated marginal means, black dots represent raw data and error bars equal 95% confidence intervals.

There was a significant effect of initial infection status, and a weak effect of heat treatment on forward-progressive motility (LRT χ^2^(1) = 4.0, *P* = 0.05 and LRT χ^2^(1) = 3.0, *P* = 0.08 respectively). There was a weak effect of initial infection status, and a significant effect of heat treatment on total motility (LRT χ^2^(1) = 3.4, *P* = 0.07 and LRT χ^2^(1) = 10.4, *P* = 0.001 respectively). The highest forward-progressive motility and total motility was in frogs initially testing negative for *Bd*, post-heat treatment. In contrast to sperm concentration results, irrespective of whether disease had been treated, frogs which initially tested negative to *Bd* had significantly higher forward progressive (OR: 2.2, 95% CIs: 1.0, 4.7) and total motility (OR: 1.8, 95% CIs: 1.0, 3.2) than frogs which tested positive. Irrespective of initial disease status, frogs which were post-heat treatment for *Bd* had significantly higher forward progressive (OR: 1.9, 95% CIs: 1.0, 3.8) and total motility (OR: 2.6, 95% CIs: 1.5, 4.5) than frogs which pre-treatment. Frogs initially testing negative for *Bd* had higher forward-progressive sperm motility both pre- (79.1%, 95% CIs: 64.9, 88.6) and post-treatment (87.7%, 95% CIs: 77.7, 93.6), compared to 63.1% (95% CIs: 49.9, 74.6) and 76.3% (95% CIs: 65.2, 84.7) respectively of frogs initially testing positive (Fig 2b). Frogs post-heat treatment for *Bd* had higher total sperm motility for both frogs initially negative (95.4%, 95% CIs: 92.2, 97.3) and positive (92.1%, 95% CIs: 88.4, 94.7), compared to 88.7% (95% CIs: 81.8, 93.1) and 81.6% (95% CIs: 74.6, 87.0) respectively in frogs pre-treatment (Fig 2c).

There was no effect of initial infection status or heat treatment on membrane integrity (LRT χ^2^(1) = 1.45, *P* = 0.23 and LRT χ^2^(1) = 0.02, *P* = 0.89 respectively). Sperm membrane integrity ranged from 95.7-97.6% across all treatment groups (Fig 2d).

### 3.4 Experiment 3: Chronic effect of heat treatment on longer-term sperm quality

In the four initially *Bd* positive frogs monitored six months post infection, there was a significant effect of time since exposure to heat treatment on sperm concentration (LRT χ^2^(2) = 24.7, *P* = 4.33×10^-6^), with decreasing sperm concentration at both the 5-week and six-month post-heat treatment timepoints (Fig 3a). Estimated marginal means (EMM) ranged from 1.6×10^8^ to 7.35×10^8^ cells/mL across all treatments. Frogs had the highest sperm concentration prior to heat treatment (EMM: 7.35×10^8^, 95% CIs: 5.8×10^8^, 9.3×10^8^). Expected sperm concentration was 3 times higher than five weeks post-heat treatment (Ratio: 2.9, 95% CIs: 2.1, 4.1) and 5 times higher than six months post-heat treatment (OR: 4.6, 95% CIs: 3.3, 6.5). Frogs six months post-heat treatment had lower sperm concentration (EMM: 1.6×10^8^, 95% CIs: 1.25×10^8^, 2.0×10^8^) than five-weeks post-heat treatment (EMM: 2.5×10^8^, 95% CIs: 2.0×10^8^, 3.2×10^8^), with sperm concentration 1.6 times higher at 5 weeks post-heat treatment compared to 6 months post-heat treatment (Ratio:1.6 95% CIs: 1.1, 2.2).

**Figure 3.**
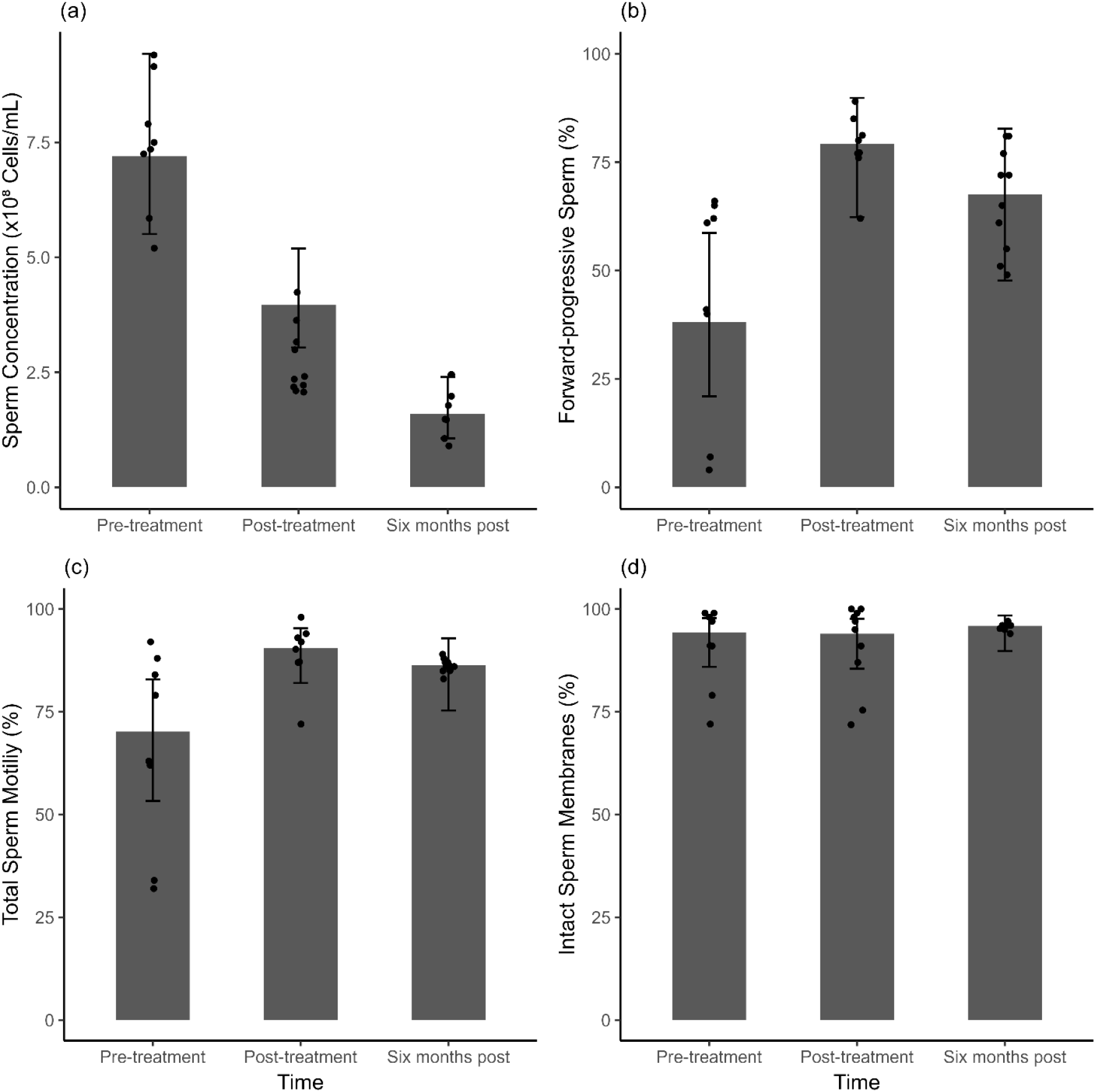
Effect of time since heat-treatment on induction of spermiation in *Ranoidea aurea* (n=4); (pre-heat treatment, five weeks post-heat treatment, and six months post-heat treatment. (a) sperm concentration (cells/mL); (b) forward progressive sperm (%); (c) total sperm motile (%), and; (d) intact membranes (%). Columns represent estimated marginal means, black dots represent raw data and error bars equal 95% confidence intervals.

There was a significant effect of time since exposure to heat treatment on forward-progressive motility (LRT χ^2^(2) = 7.1, *P* = 0.02), and on total motility (LRT χ^2^(2) = 5.8, *P* = 0.05). For both forward-progressive motility and total motility, there was an increase in sperm quality five-weeks post-heat treatment. Frogs had the lowest forward-progressive (EMM: 38.0%, 95% CIs: 21.0, 58.7) and total motility (EMM: 70.2, 95% CIs: 53.3, 82.9) counts prior to heat treatment. Frogs had the highest forward-progressive (EMM: 79.2%, 95% CIs: 62.3, 89.8) and total motility (EMM: 90.6%, 95% CIs: 82.0, 95.3) counts five-weeks post-heat treatment, with a six-fold increase in forward-progressive motility (OR: 6.2, 95% CIs: 1.9, 20.3) and four-fold in total motility (OR: 4.1, 95% CIs: 1.4, 11.5) compared to pre-heat treatment (Fig 3b and c). There was a decrease in both forward-progressive and total motility six months post-heat treatment compared to 5 weeks post heat treatment, but higher than prior to heat treatment. Forward-progressive motility was 67.6% (95% CIs: 47.7, 82.7) six months post-heat treatment. There was a 3.4-fold increase in forward-progressive motility six months post-heat treatment compared to pre-heat treatments (OR: 3.4, 95% CIs: 1.1, 11.1). Forward-progressive motility five-weeks post-heat treatment was higher than 6 months post-heat treatment, though this difference was not significant (OR: 1.8, 95% CIs: 0.6, 5.9). Total motility was 86.3% (95% CIs: 75.3, 92.9) six months post-heat treatment. There was three-fold increase in total motility six months post-heat treatment compared to pre-heat treatments (OR: 2.7, 95% CIs: 1.0, 7.4). Total motility five-weeks post-heat treatment was higher than 6 months post-heat treatment, though this difference was not significant (OR: 1.5, 95% CIs: 0.5, 4.3). There was no effect of time since exposure to heat treatment on sperm membrane integrity (LRT χ^2^(2) = 0.4, *P* = 0.83). Membrane integrity ranged from 94.0-95.9% across all treatments (Fig 3d).

### 3.5 Sperm quality from deceased frogs

To determine whether *Bd* infection induced mortality causes complete loss of gamete quality, sperm was opportunistically collected post-mortem from two frogs that died from *Bd* infection. These frogs had variable sperm quality metrics ranging in concentration from 1.39 -2.43×10^8^ cells/mL, 18.5-71.5% forward-progressive motility, 35.5 – 77.5% total motility and, 79-79.5% membrane integrity (Table 1).

**Table 1.** Sperm quality from deceased male *R. aurea* (n=2). Averages and standard deviation of two technical replicates per macerate.

|  | Sperm concentration (cells/mL)* | Forward-progressive motility (%) | Total motility (%) | Membrane integrity (%) |
| --- | --- | --- | --- | --- |
| Frog 1 | $2.43 \times 10^8 \pm 2.12 \times 10^7$ | $71.5 \pm 2.1$ | $77.5 \pm 0.7$ | $79 \pm 0$ |
| Frog 2 | $1.39 \times 10^8 \pm 4.95 \times 10^6$ | $18.5 \pm 3.5$ | $35.5 \pm 0.7$ | $79.5 \pm 0.7$ |

\* equivalent to total number of sperm recovered from testes (since recovered into 1 ml SAR).

## Discussion

This study examined how infection with *Bd* and subsequent thermal treatment influenced sperm quality and reproductive investment in male *Ranoidea aurea*. Frogs exposed to *Bd* exhibited variable infection loads and reproductive responses, with moderate infection associated with increased sperm concentration, while severe infection led to pronounced declines in both sperm concentration and motility. Thermal treatment successfully cleared the infection but reduced sperm quality across all individuals, indicating that the process of disease clearance imposed additional physiological costs. Six months after treatment, sperm motility and membrane integrity showed partial recovery, but sperm concentration remained significantly reduced, suggesting lasting impairment to spermatogenic function rather than transient suppression. These findings reveal that infection and thermal stress may interact to shape reproductive outcomes, underscoring the need to consider both disease and temperature dynamics in amphibian management and conservation.

The observed patterns in this study align with predictions from life-history theory, particularly the terminal investment hypothesis, which proposes that organisms facing elevated mortality risk will increase immediate reproductive effort to maximize fitness before death (24, 25). This study adds to the limited literature testing this hypothesis in amphibians infected with *Bd*. The increase in sperm concentration among moderately infected frogs suggests an upregulation of reproductive effort consistent with terminal investment as an acute response to infection. However, as infection severity intensified, sperm concentration and motility declined, indicating energetic trade-offs between immune defense and reproductive investment. The persistent reduction in sperm concentration post-treatment further supports the hypothesis that the energetic costs of immune activation and thermal stress may produce enduring reproductive deficits that overwhelm the capacity of an organism to generate a terminal investment. Similar context-dependent responses have been reported in other amphibians, where *Bd* infection increased reproductive output in some species but reduced it in others (12, 29). These differences likely reflect variation in infection severity, life stage, and exposure history, consistent with the dynamic threshold model proposed by Duffield et al. (26), which predicts that terminal investment occurs along a continuum shaped by survival threat and residual reproductive value.

In this study, even frogs that succumbed to *Bd* had viable and motile sperm within the testes post-mortem indicating that terminal effects of the disease are not necessarily equivalent to terminal disruption of gamete function, even if the possibility of deploying those gametes is prevented by death. While this suggests that *Bd* infection does not pose a barrier to successful reproduction in this species, there remains a paucity of data regarding the interaction of various factors, such as population wide exposure to disease, individual infection history, age and genetic diversity.

The reproductive responses observed in *Ranoidea aurea* in this study differ in both magnitude and pattern from those reported in some other *Bd*-affected amphibians. Some species have shown increased sperm production and testicular development under infection, interpreted as evidence of terminal investment (12, 29, 42). However, apparent increases in reproductive activity under mild infection may also reflect inflammatory or stress-mediated effects rather than sustained adaptive investment. Infection-induced inflammation within the testes can alter cytokine profiles, disrupt Sertoli cell function, and cause premature germ-cell release, leading to transient increases in sperm concentration or testicular activity that are not indicative of enhanced reproductive capacity (43-45). The findings in *R. aurea*, where moderate infection was associated with increased sperm concentration, but severe infection and post-treatment conditions led to marked declines, support this interpretation. This suggests that what appears as terminal investment under mild infection may instead arise from short-lived inflammatory or stress-mediated effects that precede spermatogenic exhaustion warranting further investigation. Such an acute pro-inflammatory response would only be adaptive if it could be demonstrated that fertilisation rates or offspring survival and fitness were increased.

The reduction in sperm quality observed following thermal treatment aligns with known temperature-dependent disruptions of spermatogenesis. Even modest increases in testicular temperature can impair germ-cell differentiation and reduce sperm motility and concentration through oxidative stress, mitochondrial dysfunction, and apoptosis (46, 47). Thermal exposure destabilizes the blood-testis barrier and triggers the production of reactive oxygen species, leading to germ-cell loss and DNA fragmentation. The partial recovery of motility but not concentration suggests that some aspects of sperm function may be reversible once thermally induced oxidative stress subsides, whereas damage to germ-cell populations or supporting Sertoli cells likely imposes longer-term limits on sperm production. Interestingly, membrane integrity was not significantly altered immediately following treatment, indicating that thermal and inflammatory stress primarily affected spermatogenic and maturation processes rather than post-spermatogenic membrane stability. These findings are consistent with oxidative and cytokine-mediated disruption of germ cell differentiation, rather than generalized cellular damage.

Over time, there was decreased sperm quality in frogs initially testing positive for *Bd*. The individuals initially testing negative could not be sampled at the six-month timepoint. Thus, future studies may further investigate whether long-term reproductive impacts were driven primarily by heat exposure or residual effects from infection itself. Regardless, these findings highlight that even after pathogen clearance there may be ongoing physiological consequences that reduce fertility and reproductive success in the short to medium term. Although thermal stress in this study was experimentally imposed to clear *Bd* infection and evaluate the reproductive costs of disease treatment, it also provides an important entry point for understanding how elevated temperatures associated with climate change may influence amphibian reproductive capacity. Similar physiological pathways are likely activated under natural heat extremes, suggesting that the mechanisms identified here, oxidative stress, mitochondrial dysfunction, and impaired spermatogenesis, may also operate in wild populations experiencing more frequent and intense thermal events. This overlap reinforces the value of using controlled thermal treatments not only as a disease-management tool but also as a model for assessing the broader impacts of climate warming on amphibian fertility and resilience. When viewed in the broader context of environmental change, these results suggest that the interplay between thermal stress, disease, and reproductive physiology represents a major constraint on amphibian resilience.

Evidence from hormonally induced spermiation studies indicates that sperm output can fluctuate in a sinusoidal or multi-peaked pattern across hours following induction, with concentration and motility rising and falling in recurrent waves rather than following a single peak or steady decline (48). Broader reproductive studies demonstrate that gametogenic and endocrine processes in amphibians also follow cyclical rhythms across seasonal and environmental gradients (49-51). The recognition of these temporal dynamics reinforces the need for future studies to incorporate repeated sampling across hours, days, and reproductive seasons to more accurately characterize the duration, magnitude, and variability of gametic responses to environmental and disease stress. Integrating this temporal resolution with established measures of reproductive quality will be essential for distinguishing transient stress responses from sustained shifts in reproductive allocation and for refining our understanding of amphibian resilience under changing climatic and pathogenic conditions.

## Conclusion

This study highlights how infection, temperature, and reproduction intersect to shape the physiological resilience of amphibians facing environmental change. The findings indicate that what may appear as adaptive reproductive upregulation under mild infection can instead represent short-lived physiological adjustments, and that the long-term effects of disease and recovery depend on the timing and duration of reproductive cycles. The persistence of reproductive impairment after pathogen clearance underscores that disease mitigation can itself impose physiological costs, emphasizing the importance of understanding how reproductive processes fluctuate over time. More broadly, these results point to the need for integrated research that links pathogen management, thermal tolerance, and reproductive biology, aligning with the priorities set out in the *Amphibian Conservation Action Plan* (4). By bridging these disciplines, conservation physiology can better predict how disease and climate stress jointly constrain fertility, adaptation, and long-term population persistence in a rapidly changing world.

